# Effects of conjugated linoleic acid on proliferation and differentiation of bovine intramuscular preadipocyte *in vitro*

**DOI:** 10.1101/2021.01.30.428933

**Authors:** Rong Wan, Qingxiang Meng, Zhou Zhenming, Wu hao

**Author notes:** Corresponding author. *E-mail address:* (Rong Wan).

## Abstract

Conjugated linoleic acid (CLA), a mixture of isomers of linoleic acid, has previously been shown to be able to increase intramuscular fat content *in vivo* and stimulate adipogenesis in intramuscular preadipocytes in *vitro* in pig. Unfortunately, there is little data to evaluate the effect of CLA on proliferation and differentiation of bovine intramuscular preadipocytes. This study investigated the regulation by CLA in proliferation and differentiation of bovine intramuscular preadipocytes. The results demonstrated that CLA significantly induced the expression of PPARγ and C/EBPα mRNA of bovine intramuscular preadipocytes as well as the accumulation of lipid in cultured intramuscular preadipocytes. Additionally, CLA significantly decreased the cell proportion of phase G0/G1, and remarkably increased the proportion of phase S+G2/M. Collectively, these results suggest that CLA promotes bovine intramuscular preadipocyte proliferation and differentiation.

## 1. Introduction

Intramuscular fat (Marbling) is indeed a true adipose tissue embedded within a connective tissue matrix in close proximity to a blood capillary network, primarily stored within the intramuscular adipocytes. It is one of the main factors used to determine beef quality grade in the United States (USDA, 1997), Japan (JMGA, 1988), and Korea. Insufficient intramuscular adipose and excessive subcutaneous adipose are significant beef quality challenges (Smith et al., 2006; Barnes et al., 2012). Regarding the marbling adipogenesis in the beef, the major processes involved are the proliferation of preadipocytes and their differentiation into mature adipocytes.

Conjugated linoleic acid (CLA) is a collective term for a group of octadecadienoic acids that are geometric and positional conjugated isomers of linoleic acid (Pariza et al., 2001). It is naturally generated in the rumen of ruminant animals by fermentative bacteria, which isomerizes linoleic acid into CLA. Ruminants also synthesize CLA via □9-desaturase of trans-11 octadecanoic acid (Griinari et al., 2000; Gruffat et al., 2008). It has been clearly demonstrated that treatment with CLA isomers decreases the proliferation and differentiation of murine and human preadipocytes *in vitro* (Brodie et al., 1999; Evans et al., 2000; Satory and Smith, 1999; Brown et al., 2001; Brown et al, 2003; Yeganeh et al., 2016; Yeganeh et al., 2017). However, CLA has been reported to increase intramuscular fat deposition while decrease subcutaneous fat in pigs (Dugan et al., 1997; Wiegand et al., 2002; Zhou et al., 2007). But CLA fails to inhibit the proliferation and differentiation of pig preadipocyte(Ding et al., 2000; McNeel and Mersmann, 2003). Additionally, Schmidt et al. (Schmidt et al., 2016) found that cultured adipocytes derived from dairy cows respond differently to CLA than those derived from monogastric species. These effects appear to be mediated by 2 isomers of CLA, and the 2 biologically active isomers are the CLA *cis-9, trans-11* and the CLA *trans-10, cis-12.* However, little data is available regarding the regulation of bovine intramuscular preadipocyte proliferation and differentiation by CLA. The purpose of this study, therefore, is to examine the effect of CLA on proliferation and differentiation of intramuscular preadipocytes from Chinese Luxi steers.

## 2. Materials and methods

### 2.1. Materials

Dulbecco’s modified Eagle’s medium (DMEM) and Dulbecco’s Phosphate-Buffered Saline (DPBS) were purchased from Gibco (Grand Island, N.Y.). Bovine insulin, dexamethasone, Collagenase, HEPES (powder), and conjugated linoleic acid [a mixture of CLA isomers consisting of *cis*-9, *trans-11* (50%), *cis*-12, *trans*-10 (40%), and *cis*-10, *cis*-12 (10%)] were bought from Sigma-Aldrich Fine Chemical (St. Louis, MO, USA). Fetal bovine serum (FBS) was a product of PAA Company (Austria). All other compounds were from Beijing Chemical Reagent Company (China).

### 2.2. Animal

All procedures involving animals were conducted under the approval of the Baise University, College of Agriculture and Food Engineering Animal Care and Use Committee. Three Luxi Yellow steers (18 months of age) were used in this study and approved by AEC. The cattle were fed at the National Institute of Animal Industry.

### 2.3. Cell isolation and culture

Details of tissue collection, homologous preadipocyte preparation, and the bovine intramuscular preadipocyte culturing were described as previously outlined (Aso et al., 1995). All material preparations were performed under aseptic condition, and the whole procedure was operated between the sixth and ninth passage, and the cell was cultured in a 5% CO_2_ humidified chamber at 37□.

### 2.4. Assessment of cell proliferation

Bovine intramuscular preadipocytes treated with CLA were analyzed by flow cytometry. Briefly, cells were seeded in T-25 flask at the density of 10^4^ cells/cm^2^ in DMEM containing 100 U/mL penicillin, 100 μg/mL streptomycin, 15 mM HEPES and 10% FBS (growth medium) supplemented with CLA at the concentration of 0 μM, 50 μM, 100 μM, and 150 μM. Media were changed every 2 days. 4 days later, the treated cells were collected and suspended in 15-mL sterile tubes, and then counted in hemocytometer to determine cell number. Subsequently, the cells were permeabilized with 0.1% Triton X-100 in DPBS for 30 min at room temperature.

Permeabilized cells were treated with 40 μg of RNAse/mL and then the DNA was immediately labeled with 100 μg/mL propidium iodide (PI)/mL before analysis.

### 2.5. Oil red O staining

Cells were stained with oil red O to make accumulated triacylglycerols visible. Briefly, cells were rinsed three times in DPBS and then fixed in 10% (v/v) formaldehyde for 1 h. Subsequently, the fixed cells were rapidly rinsed with isopropanol. In the end, 0.5% oil red O in isopropanol was added to the cells for 1 h, finally lipid droplets were stained with red color so that they could be visible.

### 2.6. Assay of preadipocyte differentiation

When bovine intramuscular preadipocytes were confluent (referred to day 0), the cells would be further cultured for 2 days in DMEM with mixture of 10 μg/mL insulin, 0.25 μmol/L dexamethasone, and 5% FBS (differentiation medium), and subsequently in fresh differentiation medium at different concentrations of CLA(0 μM, 50 μM, 100 μM, and 150 μM). At day 6, intracytoplasmic lipid content was determined using modified Ramirez-Zacarias’ method (Ramirez-Zacarias et al., 1992). 1.5 mL of isopropanol was added to the stained 24-well culture plates to identify the extent of preadipocyte differentiation,.Oil red O stained intracytoplasmic lipid was quantified by measuring its absorbance at 510nm on spectrophotometer.

In addition, this study was designed to investigate the regulation of individual medium components to bovine intramuscular preadipocyte differentiation. Treatment medium was consisted of base medium supplemented with: 10 μg/mL insulin, 100μM CLA, 5% FBS (DMT 1); 0.25 μmol/L dexamethasone, 100μM CLA, 5% FBS (DMT 2); 10 μg/mL insulin, 0.25 μmol/L dexamethasone, 5% FBS (DMT 3); 10 μg/mL insulin, 0.25 μmol/L dexamethasone, 100μM CLA, 5% FBS (DMT4). The amount of accumulated intracytoplasmical lipid was measured after 6-day differentiation.

### 2.7. RNA isolation and reverse transcription-polymerase chain reaction (RT-PCR)

Briefly, the housekeeping gene GAPDH was used as internal control for the identification of targeted mRNA levels. PCR was performed in 1 cycle for 5 min at 94□, several cycles of denaturation for 30 s at 94□ (details below), annealing at decreasing temperatures with increasing cycle numbers for 30 s and extension for 1 min (PPARγ, GAPDH) or 40 s (C/EBPα) at 72□. The last extension step was performed for 10 min at 72□. The primers sequences, annealing temperatures, products sizes and cycle numbers were as follows: (1) For PPARγ, forward: 5’-CAGAGATGCCGTTTTGGC-3’, reverse: 5’-AGCCGGGGATATTCTTGG-3’, annealing at 5M, product size 900 bp and 35 cycles. (2) For C/EBPα, forward: 5’-TGGACAAGAACAGCAACGAG-3’, reverse: 5’-TTGTCACTGGTCAGCTCCAG-3’, annealing at 58□, product size 130 bp and 35 cycles. (3) For GAPDH, forward: 5’-ATGCTGGTGCTGAGTATGTG-3’, reverse: 5’-GTGTCGCTGTTGAAGTCG-3’, annealing at 60□, product size 604 bp and 25 cycles. Annealing temperature and cycle numbers were determined by amplification kinetics. Gel electrophoresis with 1% (PPARγ, GAPDH) or 2% (C/EBPα) agarose gels was used for PCR products, then stained with ethidium bromide and visualized by UV-transillumination.

### 2.8. Statistical analysis

Data was expressed as least squares means ± standard error of the mean (SEM), with at least three repeats in each experimental group. Three experimental repeats were performed using BIPs from each animal sample. Results of cell proliferation and differentiation were analyzed in 1-way analysis of variance (ANOVA) with CLA treatment as the primary analysis factor. And results of various media treatment and RT-PCR were still analyzed in 1-way analysis of variance (ANOVA). Least squares means ± standard error of the mean (SEM) was used to display the study result. All data was analyzed by using SAS (version 8.0, SAS Institute, Cary, NC). Difference was considered as statistically significant at P < 0.05.

## 3. Results

### 3.1. Effect of CLA on proliferation of bovine intramuscular preadipocytes

To investigate the effect of CLA on the proliferation of bovine intramuscular preadipocytes, the cells were treated with CLA for 4 days, and then collected for analysis by flow cytometry (Fig. 2). The cell proportion of phase G0/G1 decreased significantly (P<0.05) by 9.2% in 100 μM CLA (Fig. 1; Table 1), and the proportion of phase S+G2/M remarkably (P<0.05) increased by 18.1% compared with the control group (0 μM CLA) (Fig. 1; Table 1). However, concentrations of 50 μM or 150 μM CLA supplement had no significant effect on bovine intramuscular preadipocyte proportion of G0/G1, S, and G2/M phase (Fig. 1; Table 1).

**Fig.1.**
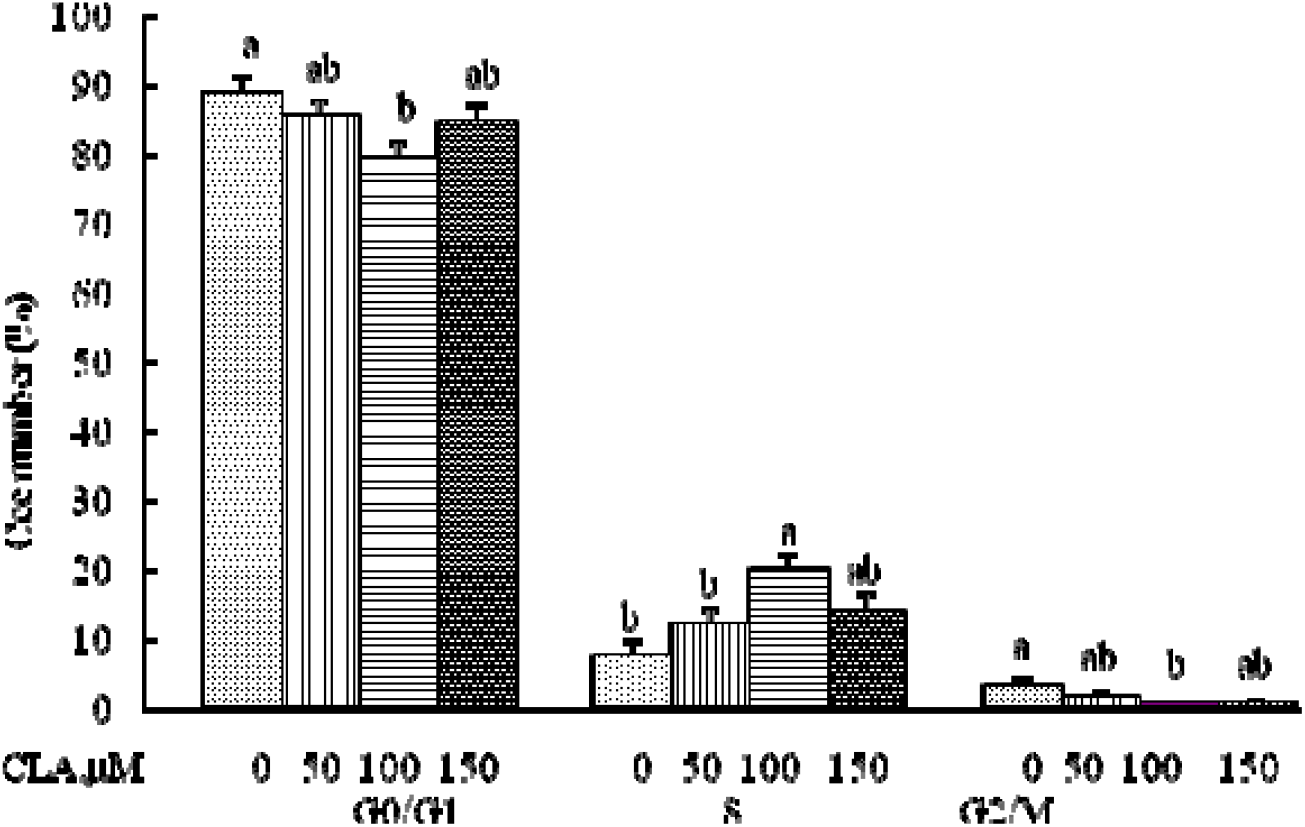
Effect of CLA on the proliferation of bovine intramuscular preadipocytes. The cell cycle is generally considered to be composed of four phases, namely the gap prior to DNA replication (G1), the synthetic phase (S), the gap after DNA (G2) and mitosis (M), and Quiescent or noncycling cells would be considered in G0 (Hartwell et al., 1989). The data was displayed as least squares means ± SEM (n= 3).

**Fig.2.**
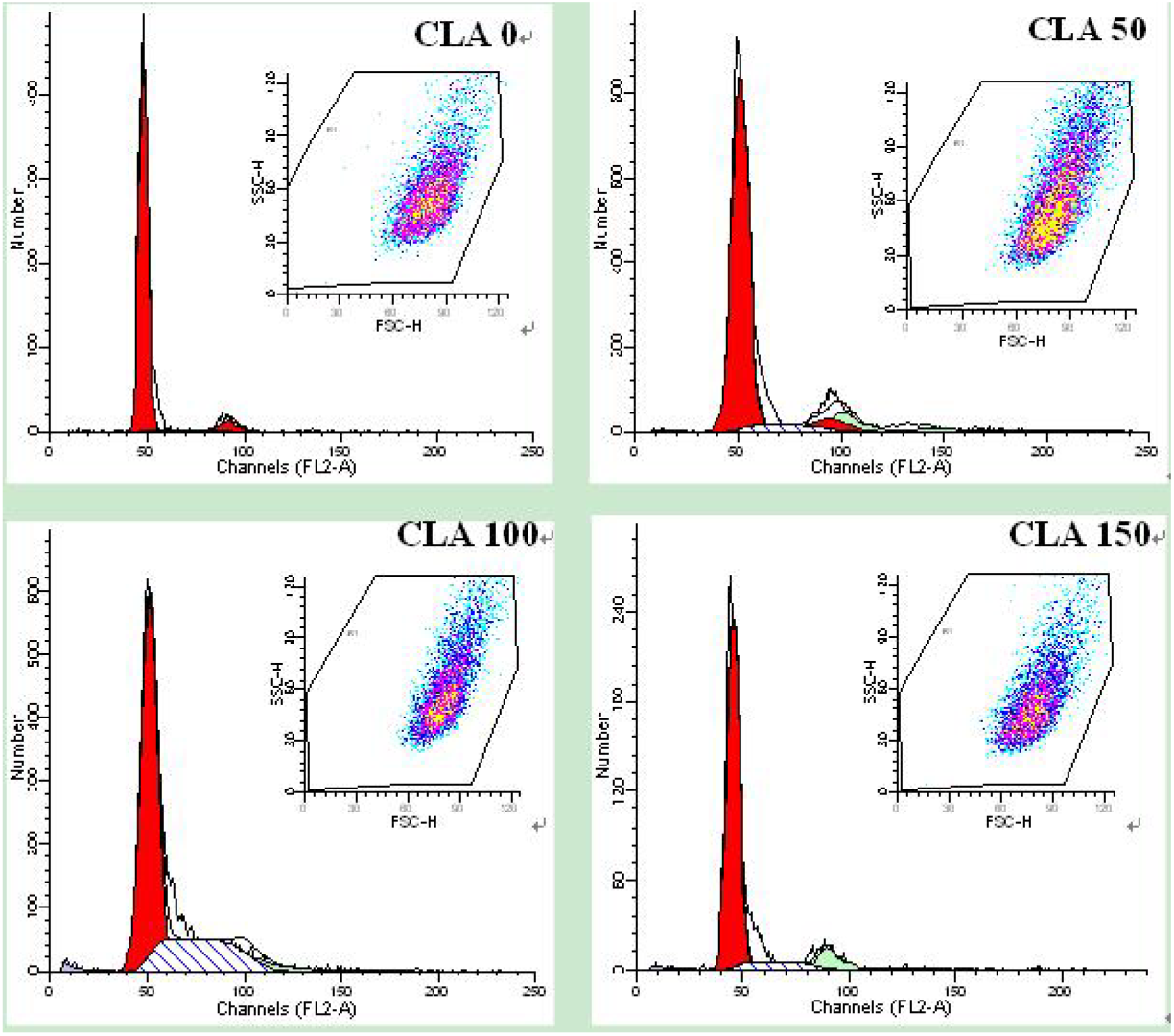
The results of the proliferation of bovine intramuscular preadipocytes were analyzed by flow cytometry.

**Table 1.**
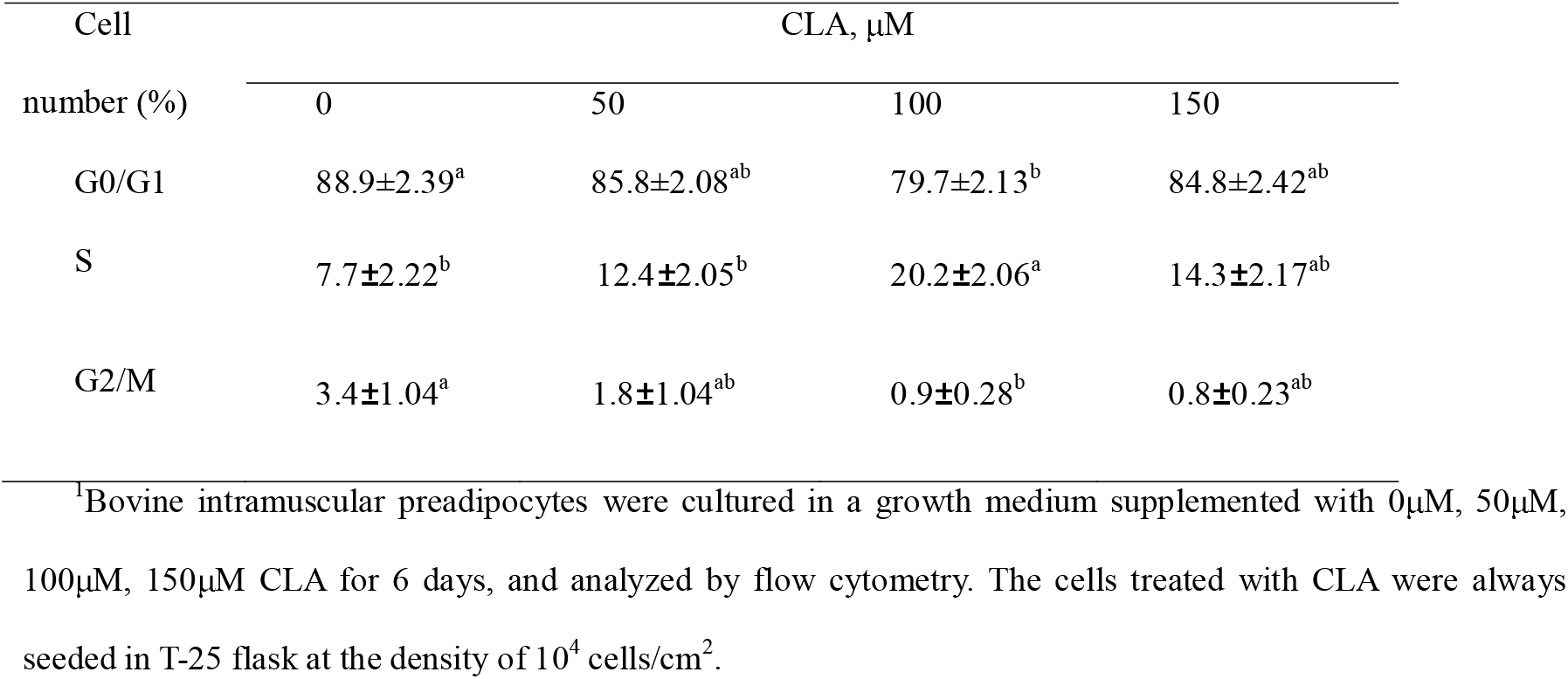
Effect of CLA on the proliferation of bovine intramuscular^1^

### 3.2. Cell morphology and lipid accumulation

To investigate the effect of CLA on the differentiation of bovine intramuscular preadipocytes, intracytoplasmic fat content in the cell which was treated with CLA for 6 days, was determined through measuring intracellular lipid droplets stained with oil red O. After the cells were treated with CLA, significant accumulation of intracellular lipid droplets was observed at day 6 (Fig. 3), and intracellular lipid content tended to increase progressively (Fig. 3). in treatments of 50, 100 and 150 μM CLA compared with control group. In addition, the results showed that CLA increased the cytoplasmic fat content of intramuscular preadipocytes in a dose-dependent manner, 1.5, 2.5, and 4 times that of the control group respectively (Fig. 4; Table 2).

**Fig.3.**
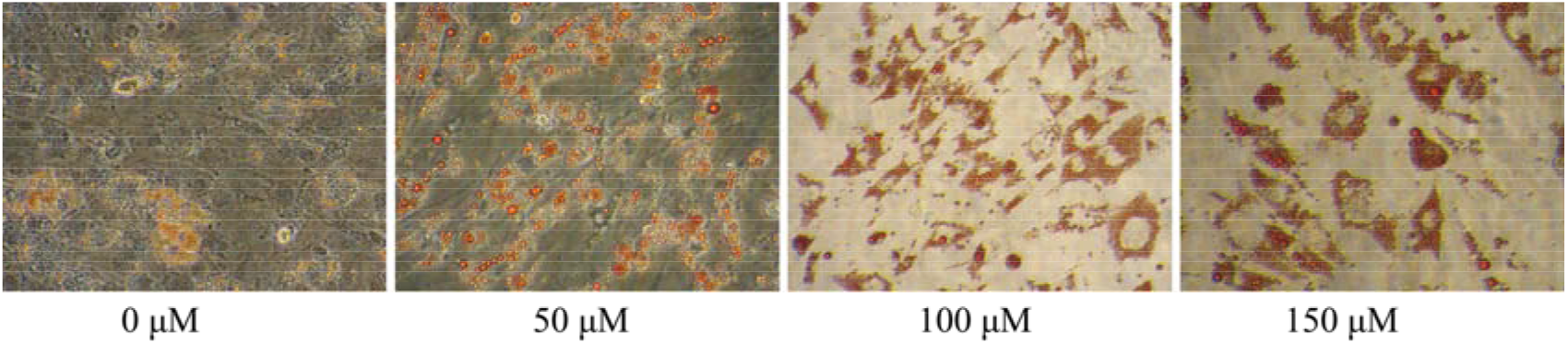
Morphological changes of bovine intramuscular preadipocyte differentiation stimulated by CLA. Intracellular lipid droplet was visualized by oil red O staining on day 6. Confluent bovine intramuscular preadipocytes were cultured in differentiation medium with the mix of 0μM, 50μM, 100μM, and 150μM CLA. Bar= 100 μm.

**Fig.4.**
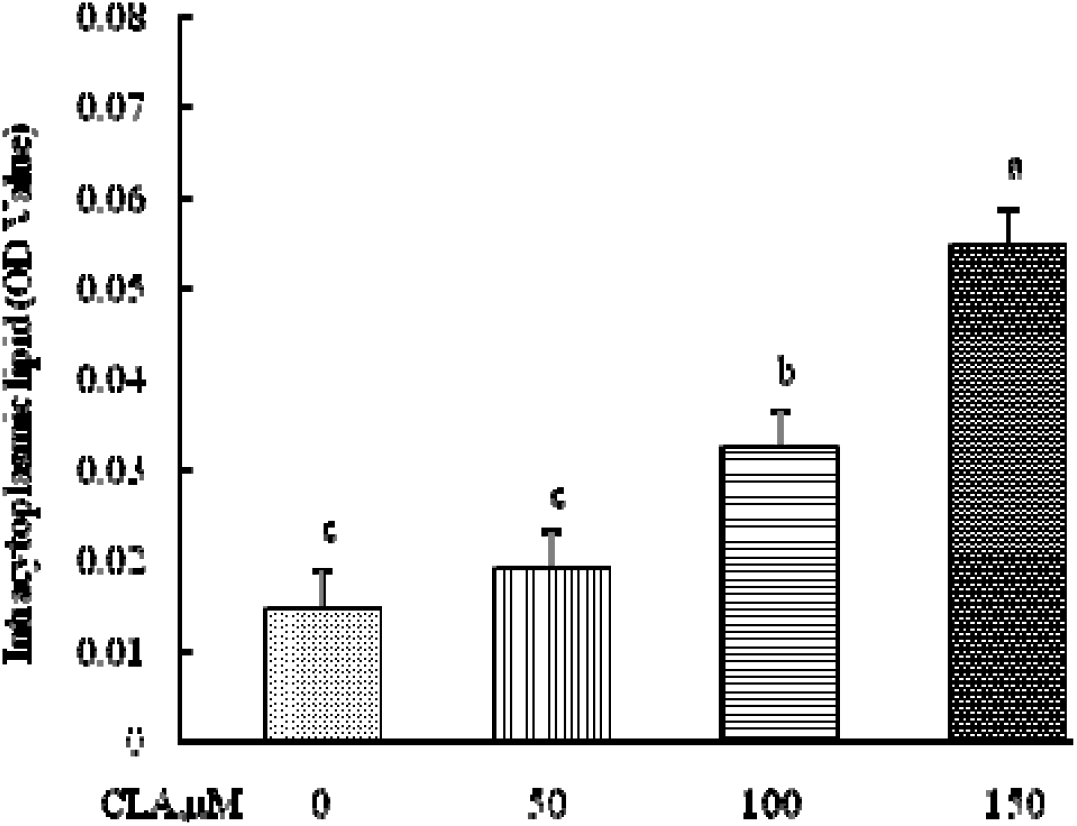
Effect of CLA on accumulation of intracytoplasmic lipid in bovine intramuscular preadipocytes. The cells treated with CLA were always seeded in 24-well plates at the density of 10^4^ cells/cm^2^, and OD Value was used to collect the number of cells per well. The data was displayed as least squares means ± SEM (n= 3). ^a, b, c^ Least squares means without a common superscript letter differ (P < 0.05).

**Table 2.** Effect of dose of CLA on the accumulation of intracytoplasmic lipid in bovine intramuscular preadipocytes^1^

| Item | CLA, $\mu\text{M}$ | | | |
| --- | --- | --- | --- | --- |
|  | 0 | 50 | 100 | 150 |
| OD Value | 0.015±0.004<sup>c</sup> | 0.019±0.004<sup>c</sup> | 0.032±0.004<sup>b</sup> | 0.055±0.004<sup>a</sup> |

<sup>1</sup> Bovine intramuscular preadipocytes were cultured in a differentiation medium supplemented with 0 $\mu\text{M}$ , 50 $\mu\text{M}$ , 100 $\mu\text{M}$ , 150 $\mu\text{M}$ CLA for 6 days, and OD Value was determined to collect the number of cells per well. The cells treated with CLA were always seeded in 24-well plates at the density of 10<sup>4</sup> cells/cm<sup>2</sup>.

### 3.3. Effect of various media components on the accumulation of intracytoplasmic lipid in bovine intramuscular preadipocytes

To determine the effects of various media components on the differentiation of bovine intramuscular preadipocyte, intracytoplasmic lipid content of the treated bovine intramuscular preadipocytes was measured. The combination of CLA, insulin, and dexamethasone resulted in the greatest intracytoplasmic lipid content (DMT4, Fig. 5), whereas the removal of dexamethasone, insulin or CLA reduced intracytoplasmic lipid content by 74%, 68% and 31% respectively (DMT1, DMT2, DMT3, Fig. 5) compared with DMT4 group (P < 0.05).

**Fig.5.**
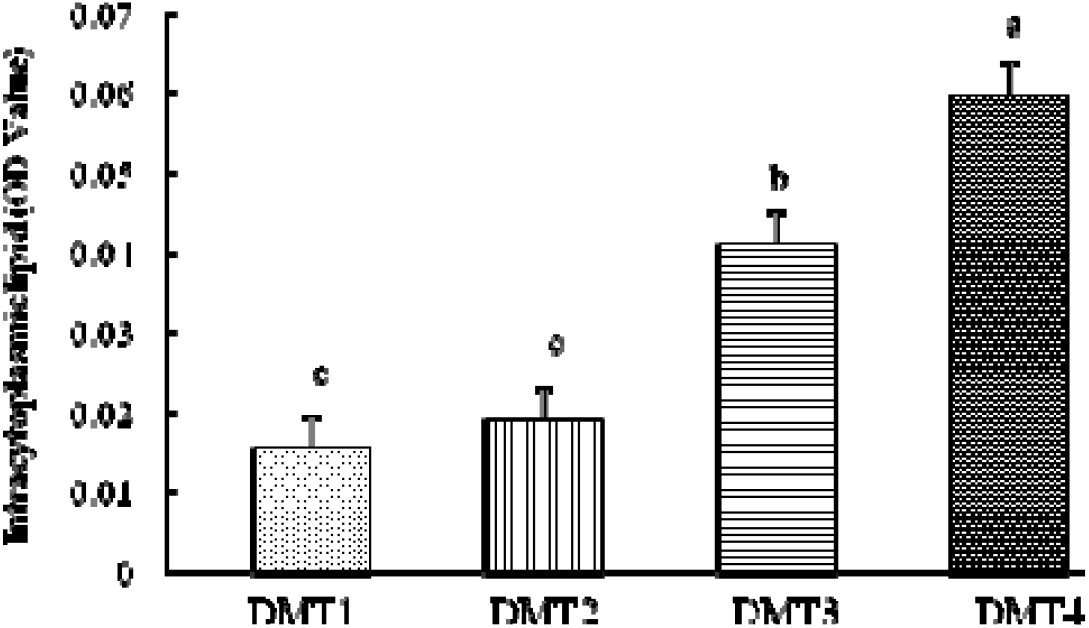
Effect of various media components on the accumulation of intracytoplasmic lipid in bovine intramuscular preadipocytes. The treatment of various media components included (1) 10 μg/mL insulin, 100μM CLA, 5% FBS (DMT 1);(2) 0.25 μmol/L dexamethasone, 100μM CLA, 5% FBS (DMT 2); (3) 10 μg/mL insulin, 0.25 μmol/L dexamethasone, 5% FBS (DMT 3); (4) 10 μg/mL insulin, 0.25 μmol/L dexamethasone, 100μM CLA, 5% FBS (DMT4). The cells treated with CLA were always seeded in 24-well plates at the density of 10^4^ cells/cm2, and OD Value was used to collect the number of cells per well. The data are expressed as least squares means ± SEM (n= 3). ^a, b, c^ Least squares means without a common superscript letter differ (P<0.05).

### 3.4. mRNA expression of adipogenic marker genes during adipogenesis of bovine intramuscular preadipocytes

To further reveal the effect of CLA on the differentiation of bovine intramuscular preadipocytes, the effect of CLA on the expression of key adipogenic transcription factors was examined. The expression of mRNA transcripts for these genes on day 6 after induction is shown in the result (Fig. 6). CLA treatment significantly up-regulated the expression of PPARγ and C/EBPα mRNA of bovine intramuscular preadipocytes compared with the control (P < 0.05, Fig. 6). Most importantly, the concentration of PPARγ mRNA increased threefold in the CLA-treated cells, whereas the concentration of C/EBPα mRNA doubled (Fig. 6). There was no difference in PPARγ and C/EBPα mRNA expression among the CLA-treated groups.

**Fig.6.**
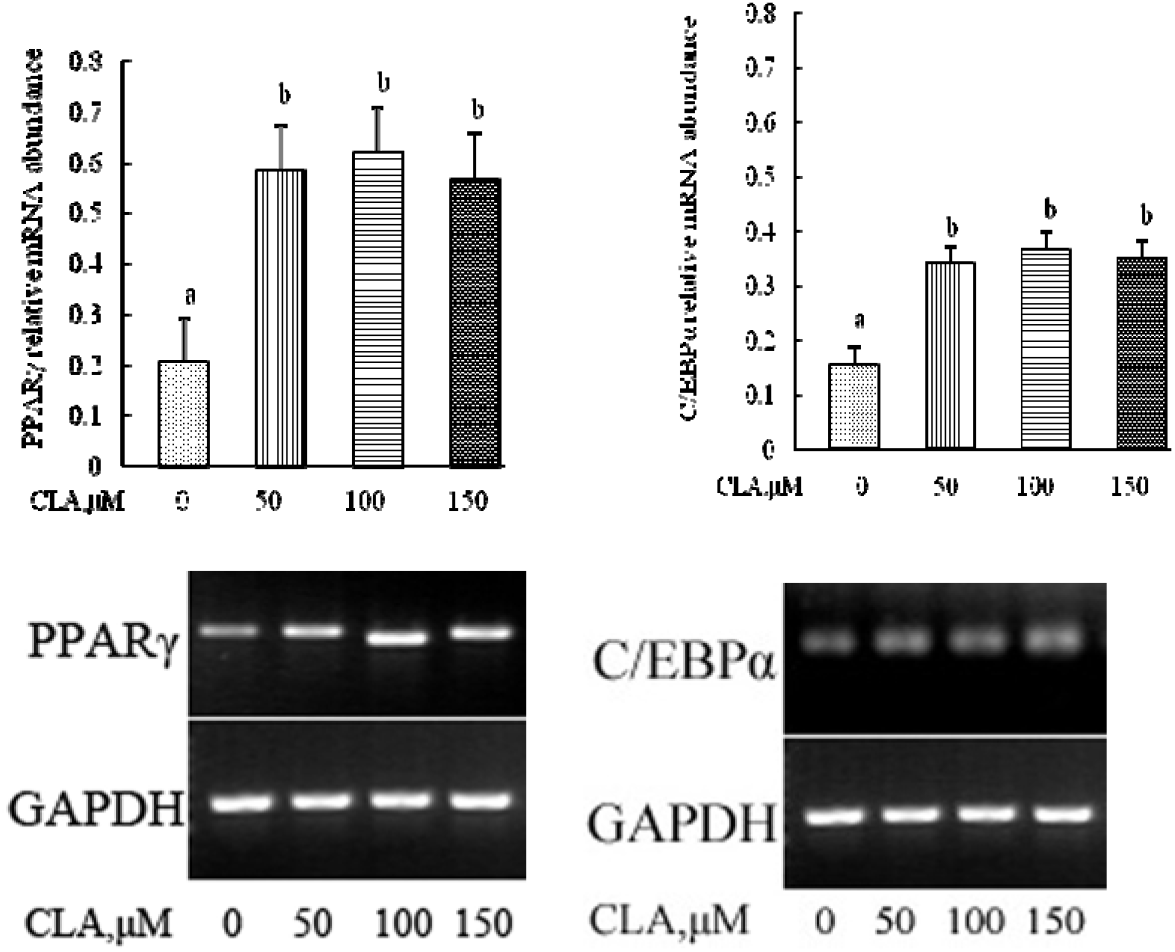
mRNA expression of marker for adipogenesis (PPAR and C/EBPα) in differentiated bovine intramuscular preadipocytes. Confluent bovine intramuscular preadipocytes were cultured in differentiation medium with the mix of 0 μM, 50μM, 100μM, and 150μM CLA. RT-PCR was carried out using gene specific primers. The PCR products were run on 1% (PPARγ, GAPDH) or 2% (C/EBPα) agarose gels, stained with ethidium bromide and visualized by UV-transillumination. The expression of PPARγ and C/EBPα were normalized by the expression of GAPDH. The data are expressed as least squares means ± SEM (n= 3). ^a, b^Least squares means without a common superscript letter differ (P < 0.05).

## 4. Discussion

This study documented the effect of CLA on the proliferation and differentiation of bovine intramuscular preadipocytes *in vitro.* Several studies have reported that CLA is an effective regulator of body fat accumulation and retention (Pariza et al., 1996; Park et al., 1997; Belury and Kempa-Steczko, 1997; Sisk et al., 1998; Azain, 2003; Dugan et al., 2004; Yeganeh et al., 2017). Satory and Smith (Satory and Smith, 1999) found that low levels (1-6 μM) of crude mixture of CLA isomers reduced the proliferation of 3T3-L1 preadipocytes. Contrary to these data, Brodie et al. (Brodie et al., 1999) found that 25-100 μM of a crude mixture of CLA isomers inhibited the proliferation of cultures of 3T3-L1 preadipocytes. Additionally, McNeel and Mersmann (McNeel and Mersmann, 2003) also found that 50 μM of a mixture of CLA isomers decreased porcine adipocyte growth. However, the current results clearly show that CLA at 100 μM significantly decreases the cell proportion of phase G0/G1, and remarkably increases the proportion of phase S+G2/M. Both the proportion of S and G2/M phase refers to the extent of cell proliferation. The above results demonstrate that proliferation of bovine intramuscular preadipocytes is promoted by CLA at appropriate concentration. Interestingly, previous studies indicated that CLA treatment had no effect on porcine intramuscular stromal-vascular cells (Zhou et al., 2007). The divergence of results with these experiments may result from different culture condition, species-specific response, or distinction between clonal and primary preadipocytes (stromal-vascular cells).

A novel aspect of this investigation is the observation that CLA significantly increased intracytoplasmic lipid content of bovine intramuscular preadipocytes in a dose-dependent way, visualized by oil red O staining.It is demonstrated that culture of mature murine 3T3-L1 adipocytes treated with 20-200 μM of crude mixture of CLA isomers for 2 days has 8% less lipid compared with control group (Park et al.,1997).it is also reported subsequently that culture of mature 3T3-L1 adipocytes treated with 100 μM of a crude mixture of CLA isomers had 55% less triglyceride content and 1.8 fold more lipolysis than that of control group (Park et al.,1999). Additionally, several studies also demonstrate that mixed isomers of CLA reduce triglyceride content in differentiating 3T3-L1 preadipocytes (Park et al., 1999; Evans et al., 2000; Evans et al., 2001). However, differences observed among these investigations may have been due to different species, especially adipose tissue depots. It has been demonstrated that preadipocytes from different depots in rats (Sztalryd et al., 1991; Kirkland et al., 1996), porcines (Hausman and Poulos, 2004; Poulos and Hausman, 2006; Zhou et al., 2007), humans (Wabitsch et al., 1996) become varied in differentiation in vitro under identical condition.

Insulin is known to stimulate adipogenesis of swine (Suryawan et al., 1997). Exclusion of insulin from differentiation media prevented the adipogenesis of ovine preadipocytes. Combination of insulin and DEX is necessary to stimulate lipid filling of adipocytes (Adams, Flint et al., 1996, Ramsay, Rosebrough et al., 2003). Additionally, glucocorticoid is necessary for preadipocytes differentiation, e.g, porcine preadipocytes continuously exposed to a glucocorticoid undergo considerable differentiation (McNeel and Mersmann, 2003). Glucocorticoids play, at physiological concentration, a permissive role of terminal differentiation in confluent Ob1771 mouse preadipocytes (Gaillard et al., 1991) and 3T3-L1 adipocytes (Wu et al., 1996). Our current study also indicates that the combination of CLA, insulin, and dexamethasone lead to the greatest intracytoplasmic lipid accumulation (DMT4), whereas the removal of dexamethasone, insulin or CLA reduces intracytoplasmic lipid content by 74%, 68% and 31% respectively (DMT1, DMT2, DMT3) compared with DMT4 group. Thus, there are synergetic effects among DEX, insulin, and CLA in enhancing bovine intramuscular preadipocyte differentiation. Both DEX and insulin may act as modulators of preadipocyte differentiation and be essential for adipogenesis in adipocytes.

As preadipocytes differentiate, the mRNA concentration of key transcription factors, such as PPARγ and C/EBPα is expected to increase accordingly (MacDougald and Lane, 1995; Rosen et al., 2000). Adipocyte differentiation is the result of transcriptional remodeling that leads to the activation of multiple adipocyte-related genes. Transcription factors such as PPARγ and C/EBPα are considered to play a crucial role in preadipocyte differentiation and act in concert to generate fully mature adipocytes (Evan et al, 1999; Paul, 2001; Evan et al., 2002; Feve, 2005; Evan and Ormond, 2006; Masaaki et al., 2008). The results of this study are consistent with the finding that, *in vivo,* the expression of PPARγ mRNA in longissimus muscle is induced by dietary CLA treatment (Meadus et al., 2002). Our data also demonstrates that CLA treatment significantly up-regulate the expression of PPARγ mRNA of bovine intramuscular preadipocytes. The other two studies also support these findings (Zhou et al., 2007; Evans et al., 2001). Since PPARγ activity is regulated by fatty acid ligands (Spiegelman, 1998; Krey et al., 1997; Schoonjans et al., 1996), and the CLA isomers has been reported to be weak agonists of PPARγ (Clement et al., 2002; Yu et al., 2002), we speculate that the CLA isomers might act as agonists of PPARγ. Another possible target gene of CLA is C/EBPα. C/EBPα is not expressed during proliferation in 3T3-L1 preadipocytes (Umek et al., 1991) but is highly induced by the onset of differentiation in 3T3-L1 preadipocytes (Christy et al., 1991). Our results are consistent with the stimulation effect of C/EBPα gene expression by CLA treatment on 3T3-L1 preadipocytes. Consequently, our data indicates that there is clear correlation between the accumulation of intracytoplasmic lipid and the expression of PPARγ and C/EBPα mRNA.

In conclusion, our data suggests that CLA can promote bovine intramuscular preadipocyte proliferation and differentiation, which provides new perspective on the control of CLA on intramuscular preadipocyte proliferation and differentiation to improve meat quality in beef industry. However, CLA may have different effect on adipogenesis depending on isomer type and concentration, culture condition, and adipose tissue depots. Therefore, detailed in-depth research is required to verify exactly how CLA influence bovine preadipocyte proliferation and differentiation in more specifical perspective.

## Acknowledgments

The authors wish to thank Jian Ding, Xiaoling Xu and Liping Sun for their assistance during the experiments.

## Funding

This study was supported by the research funding of Baise University (DC2000002684).

## Competing Interests

The authors declare they have no conflicts of interest.

## Data and Material Availability

Data are available from the corresponding author.

## Author Contributions

Rong Wan: Conceptualization, Methodology, Validation, Formal Analysis, Investigation, Writing-Original Draft, Supervision, Project Administration; Qingxiang Meng: Conceptualization, Resources, Writing-Review & Editing; Zhou Zhenming: Conceptualization, Methodology, Writing-Review & Editing; Hao Wu: Conceptualization, Writing-Review & Editing.

